# A New aspect of the pathology of brain tuberculosis: *Mycobacterium tuberculosis* infects and alters human neural progenitor cells

**DOI:** 10.64898/2026.08.16.745136

**Authors:** Thanthrige Thiunuwan Priyathilaka, Melinda Herbath, Mohan Kumar, Collin James Laaker, Michael P. Schwartz, Connie Lebakken, Zsuzsanna Fabry, Matyas Sandor

## Abstract

Brain tuberculosis remains associated with high mortality, and many survivors exhibit cognitive impairments. Progress in understanding the disease is hindered by the lack of human models. In this study, human neural organoids were infected, revealing that a subpopulation of neural progenitor cells (NPCs) is directly infected by apoptotic cell receptors expressed by NPCs, mediating bacterial uptake. Phagocytosed bacteria were localized in late endosomes, lysosomes, and the cytoplasm. Cytoplasmic bacteria frequently formed cords, indicating limited control of bacterial expansion. Immunostaining demonstrated that infected NPCs produce a type I interferon (IFN) response, corroborated by increased expression of type I IFN and IFN-regulated genes detected by RNA sequencing. Pathways related to innate immune response, cell death, and proliferation were also activated following *Mycobacterium tuberculosis* (Mtb) uptake by NPCs. The addition of color-coded microglia and monocytes to 3D neural organoids and NPCs revealed cross-infection of NPCs and other phagocytes by Mtb, suggesting a mechanism by which NPCs may access the bacteria. Infection of NPCs resulted in increased cell death, inhibition of neural differentiation, and reduced proliferation, effects that were partially mitigated by anti-IFNα treatment. Differentiated neurons were not infected. These findings indicate that brain organoids and NPC-based in vitro platforms provide a novel approach for studying brain tuberculosis. Decreased NPC function may contribute to brain tuberculosis-induced cognitive disease.

## Introduction

Mtb primarily infects the lungs, but the bacteria can disseminate and cause disease in other organs, including the liver, bone marrow, and brain(Moule and Cirillo 2020; Kulchavenya 2014; Mehta et al. 1991). Although dissemination to the brain occurs in only a small fraction of tuberculosis patients (1-5 %), Mtb is the most common bacterial infection of the central nervous system (CNS)(Rock et al. 2008). Despite advances in the treatment of systemic tuberculosis, the mortality rate of CNS-tuberculosis (CNS-TB) remains extremely high. Approximately half of infected patients are expected to die, and half of the survivors experience severe cognitive impairments due to permanent neurological damage(Marais et al. 2010; Gerber and Nau 2010; Garg et al. 2010; Karstaedt, Valtchanova, et al. 1998; Arvanitakis et al. 1998; Karstaedt, Jones, et al. 1998; Afghani and Lieberman 1994). Despite the significant public health impact of CNS-TB, understanding of disease progression, Mtb dissemination within CNS cells, pathogenic mechanisms, cellular fate post-infection, and anti-mycobacterial immunity in the brain parenchyma remains limited. This lack of knowledge hinders the development of improved therapies for CNS-TB. The National Institutes of Health (NIH) convened a meeting to identify obstacles to more effective CNS-TB treatments and concluded that the absence of human experimental models is a major road block to progress(Jain et al. 2018). To date, most knowledge regarding the pathogenesis and molecular mechanisms of human CNS-TB is derived from postmortem human samples and animal models(Rock et al. 2008). However, as Mtb primarily affects humans, these animal models do not fully recapitulate human disease pathogenesis. Therefore, this study introduces human neural organoids as a novel in vitro platform for modeling CNS-TB.

Human brain organoids are complex, laboratory-grown three-dimensional tissues typically derived from human induced pluripotent stem cells (hiPSCs) that recapitulate features of the developing human brain(Lancaster et al. 2013). These organoids have proven to be powerful and versatile platforms for studying infectious diseases of the human CNS(Priyathilaka et al. 2022). Brain organoids have been utilized to investigate a range of human CNS infections, including Zika virus(Krenn et al. 2021; Cugola et al. 2016; Qian et al. 2016), SARS-CoV-2(McMahon et al. 2021; Jacob et al. 2020; Ramani et al. 2020), human cytomegalovirus (HCMV)(Sun et al. 2020; Sison et al. 2019; Brown et al. 2019), human herpes simplex virus-1 (HSV-1)(Krenn et al. 2021; Qiao et al. 2020; D’Aiuto et al. 2019), human immunodeficiency virus (HIV-1)(Dos Reis et al. 2020), *Toxoplasma gondii*(Seo et al. 2020), and human cerebral malaria(Harbuzariu et al. 2019). In the context of tuberculosis studies, it has been demonstrated that the primary route of brain infection is via infected blood monocytes(Gilpin et al. 2021). Mtb infection infected a subset of organoid cells suggesting the presence of cells with phagocytic capacity, even if the absence of myeloid cells in the neural organoids. The data indicate that a subpopulation of neural progenitor cells (NPCs), are susceptible to Mtb infection.

NPCs are multipotent stem cells in the CNS capable of differentiating into various cell types, including neurons, oligodendrocytes, and astrocytes(Martínez-Cerdeño and Noctor 2018). In addition to their primary functions, NPCs can act as non-professional phagocytes, and phagocytic uptake of apoptotic cells by NPCs has been previously documented(Lu et al. 2011). NPCs are critical for brain development and can differentiate into brain organoids under controlled in vitro conditions(Schwartz et al. 2015). They are abundant in the developing CNS and are also present in the subventricular zone (SVZ) of the adult brain, where they contribute to the replacement of olfactory neurons, and in the dentate gyrus (DG) of the hippocampus, where they support learning and memory(Makrygianni and Chrousos 2023; Ginisty et al. 2019; Leeson et al. 2018). Adult neurogenesis is a dynamic and tightly regulated process, and Mtb access to this system may disrupt NPC functions, potentially contributing to brain tuberculosis pathology. The present study investigates the interaction between NPCs and Mtb to elucidate how NPC function is altered by infection. Extensive analyses using three-dimensional neural organoids and two-dimensional NPC cultures demonstrate that NPCs can ingest Mtb in vitro. The study further identifies the mechanisms of Mtb uptake, the intracellular localization of internalized bacteria, the fate of infected NPCs, and the genes and molecular pathways regulated following infection. Previous modeling of other brain infections, such as Zika virus, using three dimensional (3D) brain organoids and two dimensional (2D) NPCs, has revealed broad type I interferon responses(Krenn et al. 2021; Tabari et al. 2020; Liu et al. 2019). A similar response is observed in NPCs following Mtb infection. Additionally, the study demonstrates that Mtb-infected monocytes or microglia can disseminate bacteria to NPCs. The infection of NPCs by Mtb represents a novel aspect of brain tuberculosis pathology. Given the critical role of NPCs in learning, impaired function of Mtb-infected NPCs may contribute to the cognitive deficits associated with brain tuberculosis.

## Results

### H37Rv Mtb infects the cells of hiPSC-derived neural organoids

To investigate the interaction between human brain cells and Mtb, hiPSC-derived neural organoids were infected with fluorescently labeled H37Rv Mtb-tdTomato for three days. Samples were then immunostained for the neuronal marker TUJ1 and anti nestin or SOX2 antibodies for the NPCs and infected organoids were analyzed using confocal microscopy (Fig. 1a and 1b). An association of Mtb bacilli (red) with cells of the neural organoids was observed (Fig. 1b). As these neural organoids lacked a myeloid phagocytic cell population, Mtb uptake was not anticipated. As shown in Fig. 1c and 1d, Mtb was associated with both nestin-positive (Fig. 1c) and SOX2-positive (Fig. 1d) cell populations. Flow cytometric analysis confirmed these findings, showing that 2.2% of neural organoid cells were infected with H37Rv Mtb (Fig. 1e, upper panel) and that most infected cells were negative or low for TUJ1 expression (Fig. 1e, lower panel). Collectively, these results indicate that a subpopulation of NPCs is susceptible to Mtb infection.

**Fig. 1.**
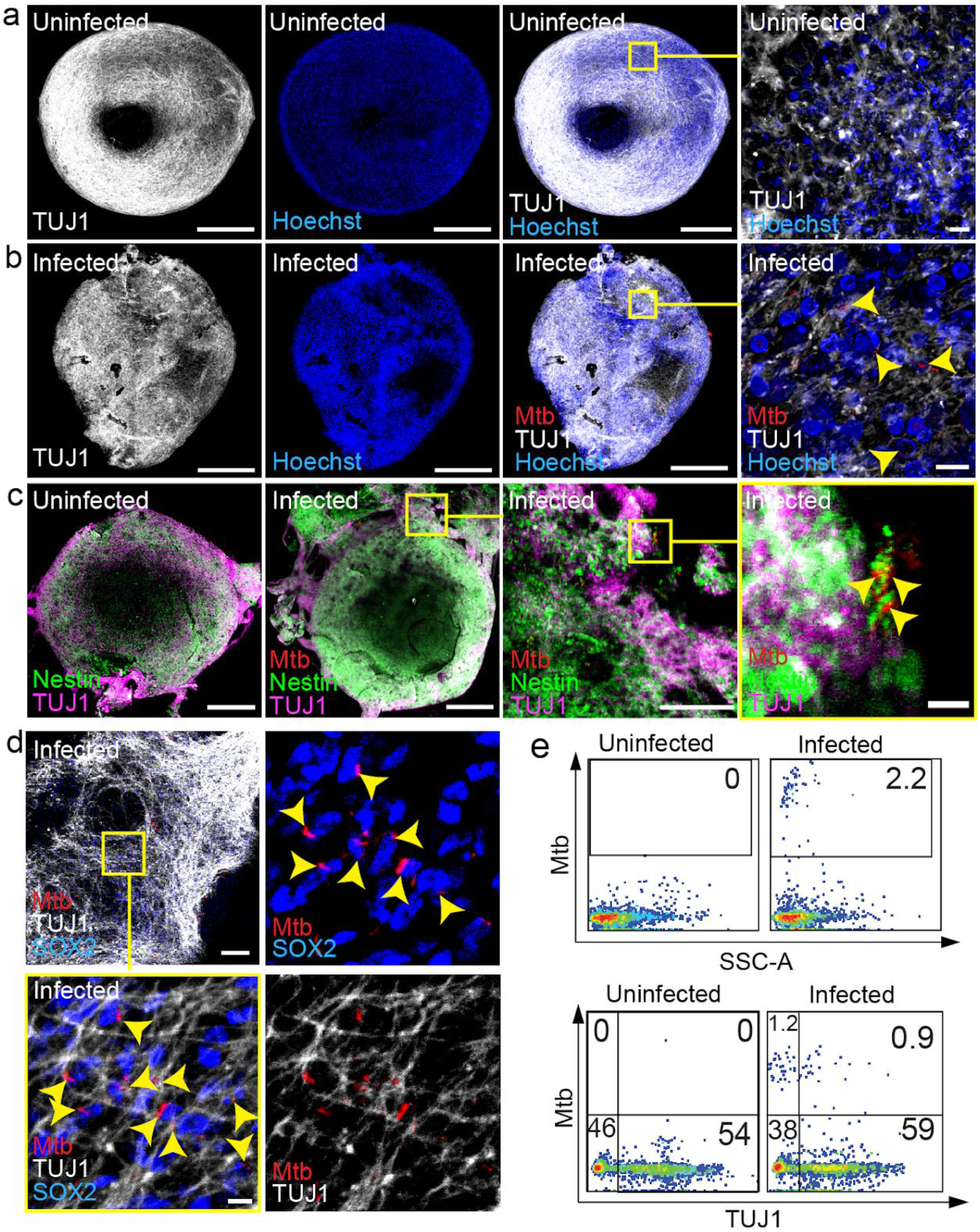
H37Rv Mtb-tdTomato (Mtb) infects human neural organoids. Human neural organoids were infected with 1 × 10^5^ CFU of H37Rv Mtb-tdTomato (red) for three days. Samples were stained for the NPC markers nestin and SOX2, as well as the neuronal marker beta III tubulin (TUJ1). Representative confocal microscopy images of uninfected **(a)** and Mtb-infected **(b)** neural organoids stained for TUJ1 (scale bar: 500 µm). High-magnification images of the boxed regions show Mtb-infected cells (yellow arrowheads) within the neural organoids (scale bar: 10 µm). **(c)** Confocal images of control (uninfected) and Mtb-infected neural organoids stained for nestin and TUJ1 demonstrate Mtb bacilli associated with nestin-positive cells (scale bars: left, 500 µm; middle, 100 µm; right, 10 µm). **(d)** Mtb-infected neural organoids show SOX2-positive cells infected with Mtb bacilli. The yellow boxed region indicates the higher magnification area, highlighting Mtb bacilli associated with SOX2-positive cells (yellow arrowheads; scale bars ;100 µm, zoom= 10 µm). **(e)** Representative flow cytometry plots of dissociated, Mtb-infected human neural organoids show the percentage of infected cells (upper panel). Flow cytometry confirms that Mtb bacilli are primarily associated with TUJ1-negative cells in neural organoids (lower panel).

### Different NPC lines are capable of uptake H37Rv Mtb and NPC can phagocytose different human Mtb strains and Mtb clinical isolates

To determine whether NPCs can internalize Mtb in vitro, NPCs derived from hiPSCs and ATCC ASC 5007 NPCs were infected with tdTomato-expressing H37Rv Mtb at a multiplicity of infection (MOI) of 1:1 for one day. Cells were subsequently labeled with the NPC marker nestin, the neuronal marker TUJ1, and the nuclear stain Hoechst. Internalization of H37Rv Mtb was visualized by confocal microscopy and quantified by flow cytometry. Both NPC lines demonstrated effective infection with H37Rv Mtb, as shown in Fig. 2a and 2e. In short-term cultures, nestin-positive and TUJ1-negative or low-expressing cells were associated with H37Rv Mtb. Confocal orthogonal projection confirmed the internalization of Mtb bacilli within NPCs (Fig. 2b). Quantitative analysis indicated that 30–40% of cells in both NPC lines were infected with H37Rv Mtb (Fig. 2d and 2e). To assess the impact of NPC differentiation on H37Rv Mtb uptake, hiPSC-derived NPCs were cultured for seven days without FGF-2 supplementation prior to infection. On day seven, cells were infected with H37Rv Mtb at an MOI of 1:1 for one day. A significantly lower percentage of H37Rv Mtb uptake was observed in these differentiated cultures compared to undifferentiated controls (Fig. 2c, 2d, and 2e), indicating that NPC differentiation reduces H37Rv Mtb uptake. To evaluate whether NPCs can internalize different Mtb strains, hiPSC-derived NPCs were infected with a human brain-derived (SK84), two sputum-derived (SK131 and SK139) Mtb clinical isolates, and the Mtb Erdman strain at an MOI of 1:1 for one day. All clinical isolates and the Erdman strain were associated with hiPSC-derived NPCs (Fig. 2f), demonstrating that both laboratory and clinical Mtb strains can infect NPCs. The SK84 brain-derived Mtb isolate exhibited higher internalization rates compared to sputum-derived isolates. Additional isolates are required to determine whether brain-derived strains demonstrate increased susceptibility to phagocytosis by NPCs. These findings provide clear evidence that NPCs can internalize Mtb in vitro.

**Fig. 2.**
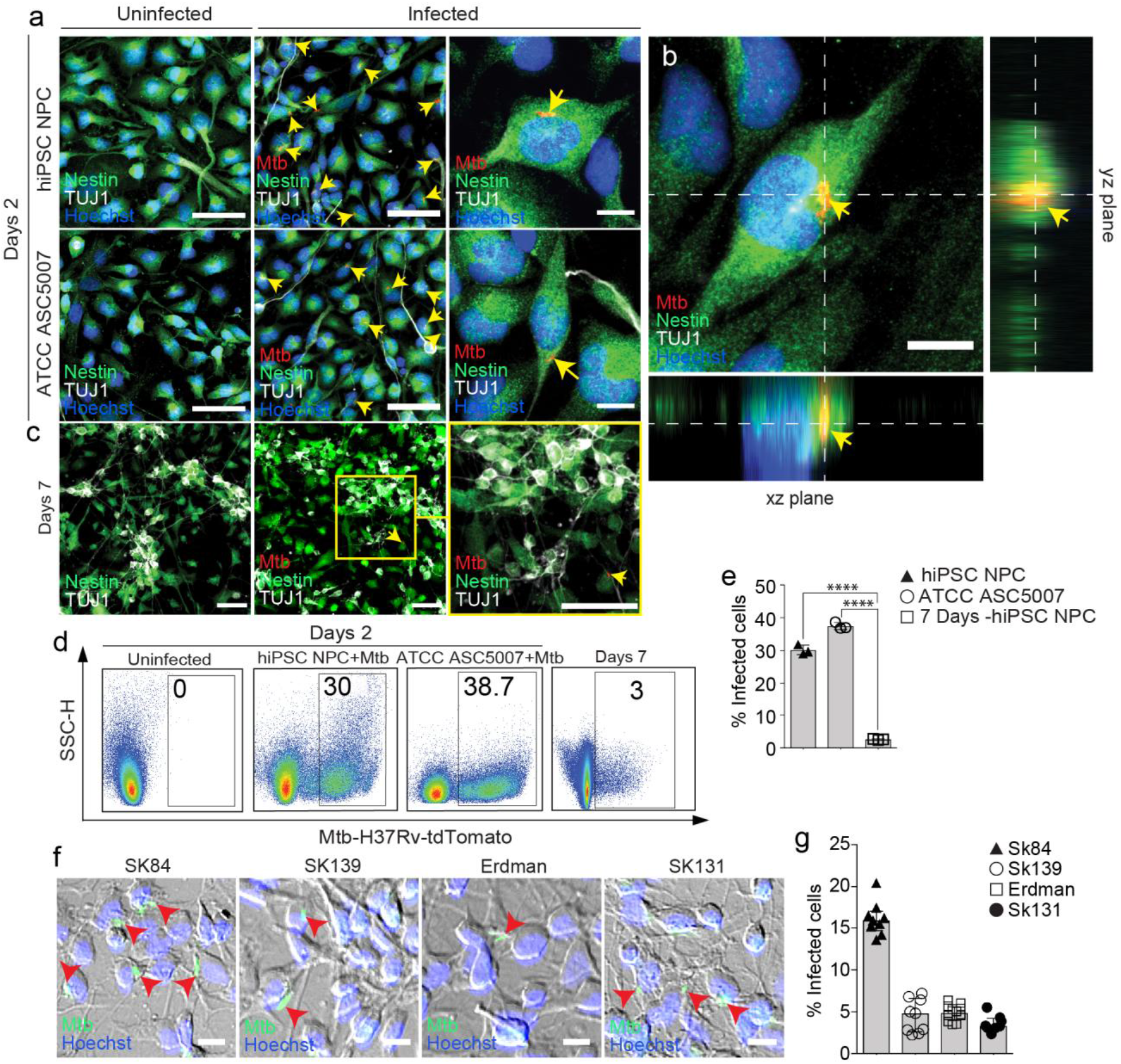
H37Rv Mtb-tdTomato infects two distinct human NPC lines, and NPC differentiation reduces Mtb uptake capacity. Undifferentiated hiPSC-derived NPCs, ATCC ACS-5007 NPCs, and partially differentiated NPCs (cultured for 7 days without FGF2) were infected with H37Rv Mtb-tdTomato (Mtb) at a multiplicity of infection of 1:1 for 1 day. **(a)** Immunocytochemical analysis of undifferentiated hiPSC-derived and ATCC ACS-5007 NPCs with or without Mtb infection. Cells were stained for the NPC marker nestin (green), neuronal marker beta III tubulin (TUJ1) (grey), and nuclear stain Hoechst 33342 (blue). Mtb-infected cells are indicated by yellow arrowheads. (Scale bars: 50 µm, 10 µm for higher magnification.) **(b)** Orthogonal projections of confocal z-stacks showing xy, yz, and xz planes of Mtb-infected NPCs. (Scale bar: 10 µm.) **(c)** Representative confocal microscopy images of infected and uninfected partially differentiated NPCs (7 days). **(d)** Representative flow cytometry plots of Mtb-infected and uninfected hiPSC-derived NPCs, ATCC ACS-5007 NPCs, and partially differentiated (7 days) NPCs. Numbers in the plots indicate the percentage of infected cells. **(e)** Frequencies of Mtb-positive cells among total cell populations. Data are presented as mean ± standard deviation from three experiments. (****p<0.0001). **(f)** Representative DIC and fluorescence microscopy images of hiPSC-derived NPCs infected with various human clinical Mtb isolates (brain: SK84; sputum: SK131 and SK139) and the Mtb Erdman strain. (Scale bars: 10 µm.) **(g)** Frequencies of hiPSC-derived NPCs positive for different human Mtb isolates and the Mtb Erdman strain.

### Several phagocytosis inhibitors partially reduce the capacity of NPCs to internalize Mtb

Several studies have suggested that NPCs exhibit phagocytic activity(Lu et al. 2011). Lu et al. (2011) demonstrated that the phagocytic activity of NPCs can be disrupted by masking phosphatidylserine (PtdSer) molecules on dying cells using annexin V(Lu et al. 2011). In the present study, it was hypothesized that the internalization of Mtb by NPCs occurs via PtdSer signals on apoptotic cells (Fig. 3a). Figure 3a illustrates the reported PtdSer receptors on NPCs: Bai1, Pros1, and purinergic P2X7 receptor (P2X7R). Each of these receptors is inhibited by Annexin V. To test this hypothesis, annexin V-treated hiPSC-derived NPCs were infected with H37Rv Mtb for one day and analyzed by flow cytometry.

**Fig. 3.**
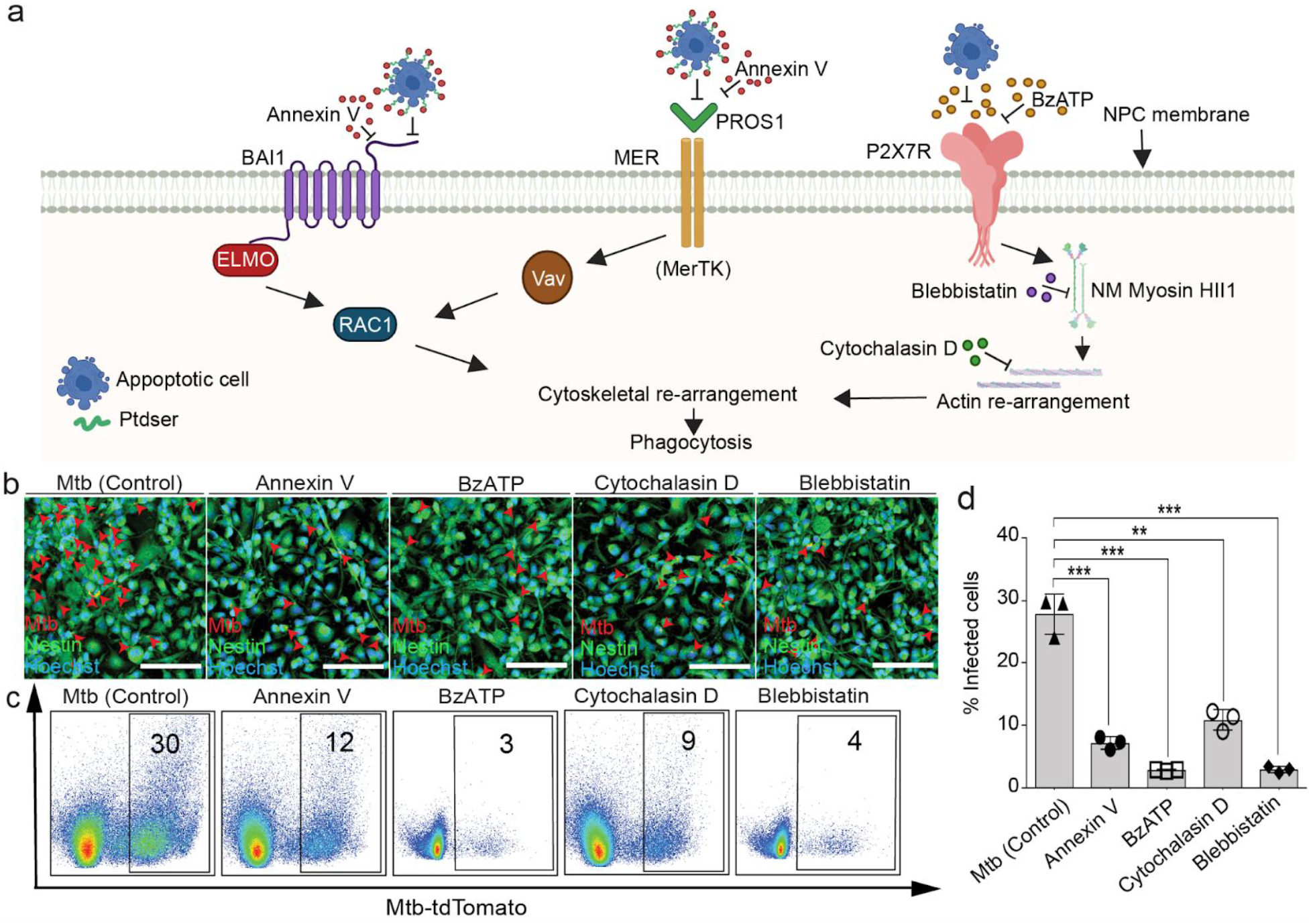
Impact of apoptotic cell phagocytosis inhibitors on Mtb uptake by NPCs. NPCs derived from hiPSCs were infected with H37Rv Mtb-tdTomato under control conditions or in the presence of 7.5 mg purified annexin V, 500 mM BzATP, 30 mM cytochalasin D, or 25 mM blebbistatin for 24 hours. Cells were subsequently analyzed using confocal microscopy and flow cytometry. **(a)** Schematic illustration of apoptotic cell uptake pathways in NPCs and their potential inhibition by the compounds used in this study. **(b)** Representative confocal microscopy images demonstrate Mtb internalization by NPCs with or without inhibitors (scale bars = 100 µm). **(c)** Representative flow cytometry plots of Mtb-infected NPCs under control and inhibitor-treated conditions, with percentages of infected cells indicated. **(d)** Frequencies of Mtb-positive cells among total cell populations in the presence or absence of inhibitors. Data are presented as mean ± standard deviation from three independent experiments. (**p<0.01, ***p<0.001).

Annexin V treatment, which inhibits these receptors, significantly reduced the uptake of H37Rv Mtb bacilli by NPCs compared to the untreated group (Fig. 3b, c, and d). Previous studies have shown that the P2X7R in NPCs can also function as a scavenging receptor for the phagocytic engulfment of apoptotic cells, latex beads, and cellular debris in the absence of their agonists(Leeson et al. 2018; Lovelace et al. 2015). P2X7R-driven phagocytosis of apoptotic cells can be inhibited by pretreatment with ATP(Lovelace et al. 2015). In the present study, to investigate the role of P2X7R in hiPSC-derived NPCs during H37Rv Mtb uptake, NPCs were infected with H37Rv Mtb in the presence of BzATP, an agonist of P2X7R. H37Rv Mtb uptake was markedly diminished following BzATP treatment compared to the untreated group (Fig. 3b, c, and d), indicating that P2X7R may play a role in H37Rv Mtb phagocytosis. Subsequently, the potential roles of cytoskeletal rearrangement and non-muscle myosin II in the cellular uptake of H37Rv Mtb-infected apoptotic cells by NPCs were examined. To this end, hiPSC-derived NPCs were infected with H37Rv Mtb in the presence of the actin polymerization inhibitor cytochalasin D and the myosin class II inhibitor blebbistatin. As shown in Fig. 3b, c, and d, treatment with cytochalasin D and blebbistatin effectively decreased the uptake of H37Rv Mtb by NPCs, suggesting that cytoskeletal rearrangement, which is essential for phagocytosis, plays a critical role in Mtb uptake. Additionally, these findings demonstrate that membrane-bound phagocytic receptors for apoptotic cells mediate the engulfment of Mtb by NPCs.

### H37Rv Mtb is present within late endosomal vesicles and is also detected in the cytoplasm of infected NPCs, where it forms intracellular cording structures

Next, the intracellular localization of Mtb following engulfment by NPCs was investigated. Transmission electron microscopy (TEM) revealed that H37Rv Mtb bacilli are associated with intracellular vesicles and are also present in the cytoplasm of NPCs one day post-infection (Fig. 4a). To confirm the intracellular localization of H37Rv Mtb, infected cells were stained for the early endosome marker Rab5A, late endosomal marker Rab7A as well as lysosomal markers LAMP-1, and Lysotracker Deep Red. Confocal microscopy demonstrated that H37Rv Mtb bacilli did not colocalize with Rab5A-positive early endosomal compartments (Fig. 4b). In contrast, most vesicle-associated H37Rv Mtb bacilli colocalized with Rab7A-positive late endosomal compartments (Fig. 4c), LAMP-1 (Fig. 4d), and Lysotracker Deep Red-positive lysosomal compartments (Fig. 4e). Confocal orthogonal projections further confirmed the internalization of H37Rv Mtb bacilli into late endosomes and lysosomes (Fig. 4c, d, and e, lower panels). Internalized Mtb inhibits the fusion of phagosomes with lysosomes to promote bacterial survival in professional phagocytes such as macrophages(Jamwal et al. 2016). Pathogenic Mtb can inhibit phagosomal maturation by manipulating host signaling mechanisms, which results in retention of Mtb within early endosomal compartments and facilitates eventual escape from phagolysosomes into the cytoplasm(Queval et al. 2017; Jamwal et al. 2016; Meena and Rajni 2010). However, this phagosomal escape mechanism has not been extensively studied in Mtb-infected non-myeloid cells. Khan et al. (2017) demonstrated that Mtb-infected mesenchymal stem cells limit intracellular bacterial growth through autophagy(Khan et al. 2017). As shown in Fig. 4g and h, cytoplasmic bacteria form intracellular cord-like structures in some H37Rv Mtb-infected NPCs three days after infection. Similarly, Mtb forms intracellular cords in lymphatic endothelial cells, which may facilitate evasion of host immunity(Lerner et al. 2020).

**Fig. 4.**
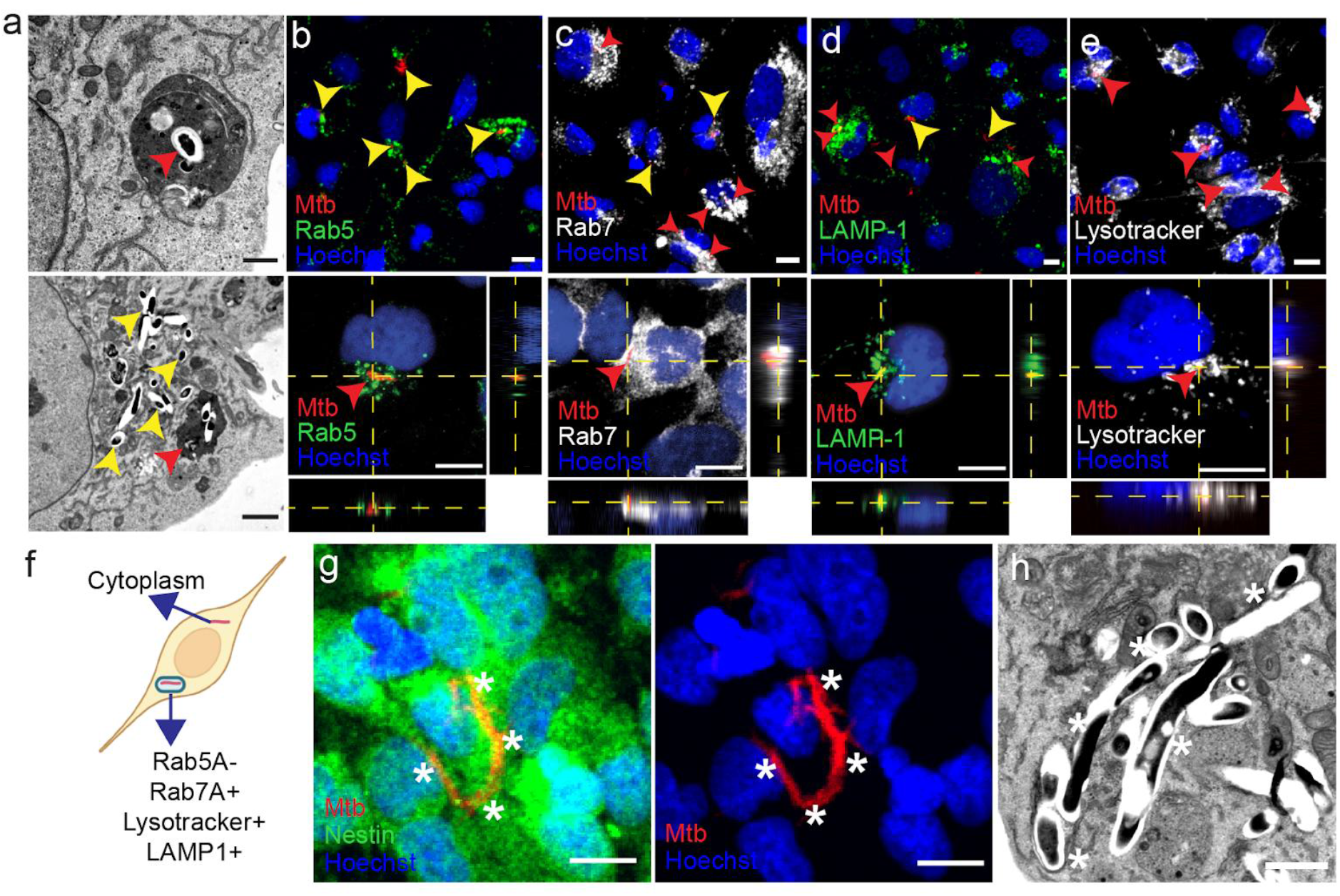
Mtb localizes within late endosomal vesicles and the cytoplasm of infected NPCs, forming intracellular Mtb cords. NPCs derived from hiPSCs were infected with H37Rv Mtb-tdTomato for either 1 or 3 days. **(a)** Representative transmission electron microscopy (TEM) images demonstrate Mtb bacilli localized within phagocytic vesicles or the cytoplasm. Mtb bacilli colocalized with phagocytic vesicles are indicated by red arrowheads, while cytoplasmic Mtb bacilli are marked in yellow (scale bar = 2 µm). Representative confocal microscopy images show that internalized Mtb does not colocalize with Rab5+ early endosomes **(b)**, but does colocalize with Rab7+ late endosomes **(c)**, LAMP-1+ **(d)**, and LysoTracker+ **(e)** phagolysosomes. Cell nuclei are stained with Hoechst 33342 (blue). Mtb (red) bacilli are indicated by red arrowheads. Scale bars = 10 µm. **(f)** Schematic representation of Mtb intracellular localization in NPCs after 1 day. Representative confocal microscopy **(g)** and TEM **(h)** images show intracellular Mtb cord formation at 3 days post-infection. Mtb cords are indicated by white asterisks.

### Alterations in Gene Expression in NPCs Following H37Rv Mtb Infection

The impact of H37Rv Mtb uptake was investigated by RNA sequencing of hiPSC-derived NPCs at 1 and 3 days post-infection (dpi) to identify regulated biological pathways. This analysis identified 283 and 479 differentially expressed genes (DEGs) in H37Rv Mtb-infected hiPSC-derived NPCs at 1 and 3 dpi, respectively. Of these, 237 genes were upregulated at 1 dpi, and 325 were upregulated at 3 dpi. To determine enriched biological processes and pathways, upregulated DEGs at both time points were analyzed using SRplot(Tang et al. 2023) and Reactome pathway analysis(Milacic et al. 2024). The top ten gene ontology (GO) terms revealed predominant enrichment of response to type I interferon (IFN) and type I IFN signaling pathways in H37Rv Mtb-infected hiPSC-derived NPCs at both 1 and 3 dpi (Fig. 5a). Additionally, type II IFN signaling and response pathways were also enriched following Mtb infection (Fig. 5a). Reactome analysis confirmed that IFN-α/β signaling were among the most prominent upregulated pathways (Fig. 5b), shown by the GO biological process findings. Defense response to viruses and related processes were also significantly enriched in H37Rv Mtb-infected hiPSC-derived NPCs at both time points (Fig. 5a). Upregulation of viral response GO terms is a characteristic feature of Mtb-infected myeloid cells. For instance, enrichment of viral response pathways has been reported in Mtb-infected human macrophage-like THP-1 cells(Wu et al. 2012) and mouse macrophages(Lee et al. 2019).

**Fig. 5.**
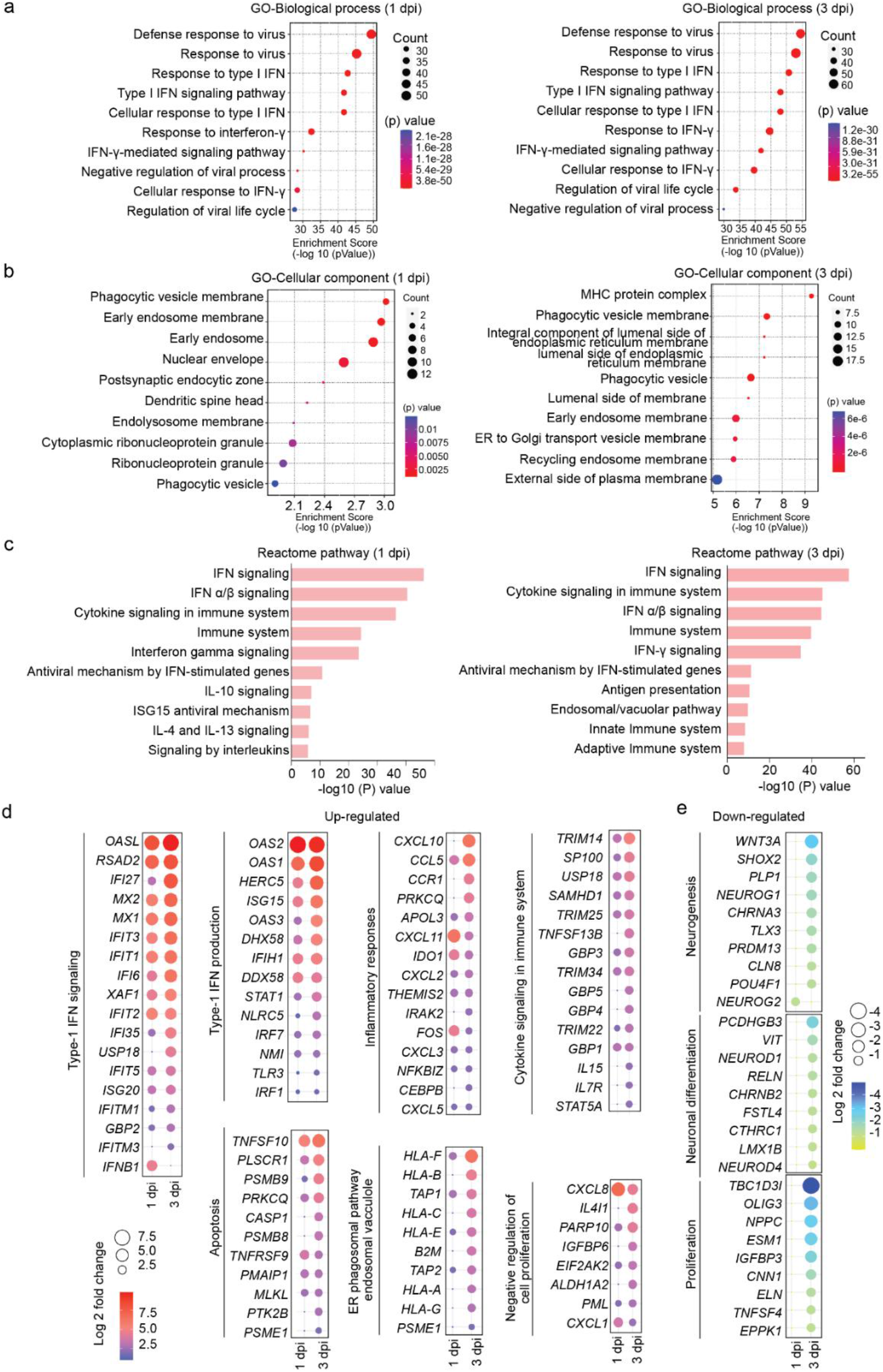
Transcriptional alterations of hiPSC-derived NPCs after Mtb infection at 1 and 3 dpi. hiPSC-derived NPCs were infected with H37Rv Mtb-tdTomato at a MOI of 1:1 for either 1 or 3 days. Total RNA was then extracted and subjected to bulk RNA sequencing. **(a)** Dot plots display the top 10 Gene Ontology (GO) biological process terms, and **(b)** cellular component terms, among up-regulated genes in NPCs at 1 and 3 days post-infection (dpi) with Mtb. **(c)** Bar graphs show the top 10 up-regulated Reactome pathway analysis hits in samples at 1 and 3 dpi. **(d)** Dot plots illustrate selected up-regulated genes, and **(e)** down-regulated genes, associated with the indicated biological processes at 1 and 3 dpi following infection.

Further analysis of the most upregulated genes demonstrated their involvement in type I IFN signaling, negative regulation of cell proliferation, type I IFN production, cell death, and cytokine signaling (Fig. 5d). Type I IFN signaling emerged as the dominant pathway compared to type II IFN signaling. Notably, several IFN-stimulated genes (ISGs), such as *OASL, RSAD2, MX1, MX2, IFIT1, IFIT2, IFIT3, IFIT5, IFI16, IFI35, XAF1, ISG20, and GBP2, were strongly induced at both 1 and 3 dpi* in hiPSC-derived NPCs (Fig. 5d). In contrast, ISGs including *IFITM1, IFITM3*, and USP18 were only induced at the later stage (3 dpi) of infection (Fig. 5d). *IFNB1* expression was particularly prominent at 1 dpi (Fig. 5d). Genes involved in type I IFN production, such as *OAS1, OAS2, OAS3, HERC5, ISG15, NLRC5*, and *IFIH1*, were upregulated at both time points (Fig. 5d). Additionally, several IFN-inducible small GTPases with established antimicrobial properties were induced following H37Rv Mtb infection. For example, guanylate-binding protein 1 (*GBP1*) confers antibacterial protection against *Mycobacterium bovis* BCG(Kim et al. 2011); *GBP2* exhibits broad antiviral activity against Zika, influenza A, and measles viruses(Braun et al. 2019); *GBP3* and *GBP5* demonstrate antiviral activity against influenza virus(Feng et al. 2017; Nordmann et al. 2012), and were detected at 1 or 3 dpi (Fig. 5d). Genes associated with inflammatory responses and cytokine signaling, including *CXCL10, CCL5, CCR1, CXCL11, IDO1, FOS*, *TRIM14, SP100*, *TRIM25, TNFSF13B, IL15,* and *IL17R*, were also induced at either 1 or 3 dpi (Fig. 5d). Furthermore, upregulation of apoptosis- and cell death-related gene signatures, such as *TNFSF10, PSMB9, CASP1, PMAIP1, FOS*, and *TNFRSF9,* was observed, with higher expression at the later stage of infection (Fig. 5d). GO cellular component analysis indicated significant enrichment of terms related to phagocytic vesicle membranes and phagocytic vesicles at 3 dpi compared to 1 dpi in H37Rv Mtb-infected hiPSC-derived NPCs (Fig. 5b). Upregulated DEGs following H37Rv Mtb infection included increased expression of *HLA-E, HLA-B*, and *HLA-C* (Fig. 5d). Previous research has shown that *HLA-E*, a non-classical class 1b MHC molecule, is highly localized to phagosomal membranes in Mtb-infected cells(Grotzke et al. 2009). Enrichment of *TAP1* and *TAP2*, recognized markers of phagosomes in H37Rv Mtb-infected cells, was also observed in hiPSC-derived NPCs in this study (Fig. 5d)(Lu et al. 2023; Harriff et al. 2013). These results collectively suggest that internalized Mtb is sequestered within the phagosomes of NPCs, even though NPCs are not considered professional phagocytes.

In contrast, the most downregulated genes were associated with neuronal functions, including regulation of membrane potential, nervous system development, neuronal differentiation, neurogenesis, and proliferation (Fig. 5e). Notably, these genes were more strongly downregulated at 3 dpi than at 1 dpi following H37Rv Mtb infection. The genes *NEUROG1*(Han et al. 2018) and *NEUROG2*(Lacomme et al. 2012), which play pivotal roles in neurogenesis and neuronal differentiation, were downregulated in hiPSC-derived NPCs upon H37Rv Mtb infection. Downregulation of *NEUROD1*, a key gene in terminal neuronal differentiation during olfactory neurogenesis(Boutin et al. 2010), was also detected in the RNASeq data of H37Rv Mtb-infected hiPSC-derived NPCs in this study. Additionally, *NEUROD4*, a critical transcription factor in neuronal development(Wang et al. 2023; Fukuoka et al. 2021) was observed to be downregulated with Mtb infection. Collectively, these data suggest that infection of hiPSC-derived NPCs with H37Rv Mtb induces host survival pathways and may serve as a valuable tool for investigating the underlying pathogenesis of CNS-TB infection and for developing potential treatments.

### H37Rv Mtb infection promotes cell death and impairs the proliferation and differentiation of hiPSC-derived NPCs in vitro

To assess the cellular effects of H37Rv Mtb infection, NPCs derived from hiPSCs were infected and incubated for 1, 3, and 5 days. Samples were stained for the proliferation marker Ki67, the NPC marker Nestin, and the neuronal marker TUJ1. Dead cells were identified using ghost-violet 450 dye. Confocal microscopy was used to acquire and quantify images. A marked reduction in Ki67-positive cells was observed in infected samples compared to uninfected controls over time, indicating that H37Rv Mtb infection impairs hiPSC-derived NPC proliferation (Fig. 6a and b). No significant increase in cell death was detected at 1 dpi relative to controls; however, a significant increase in cell death was evident at 3 and 5 dpi (Fig. 6c and d). Additionally, uninfected dead cells were found adjacent to both infected dead and live cells, suggesting that H37Rv Mtb-infected cells may induce cell death in neighboring cells through indirect mechanisms (Fig. 6c). The effect of infection on hiPSC-derived NPC differentiation was evaluated by quantifying TUJ1-positive cells over time and comparing these to uninfected samples. No significant difference in TUJ1-positive cells was observed at 1 and 3 dpi; however, a significant reduction was detected at 5 dpi in infected samples (Fig. 6d and e). These findings demonstrate that H37Rv Mtb infection induces cell death in hiPSC-derived NPCs and disrupts their proliferation and differentiation.

**Fig. 6.**
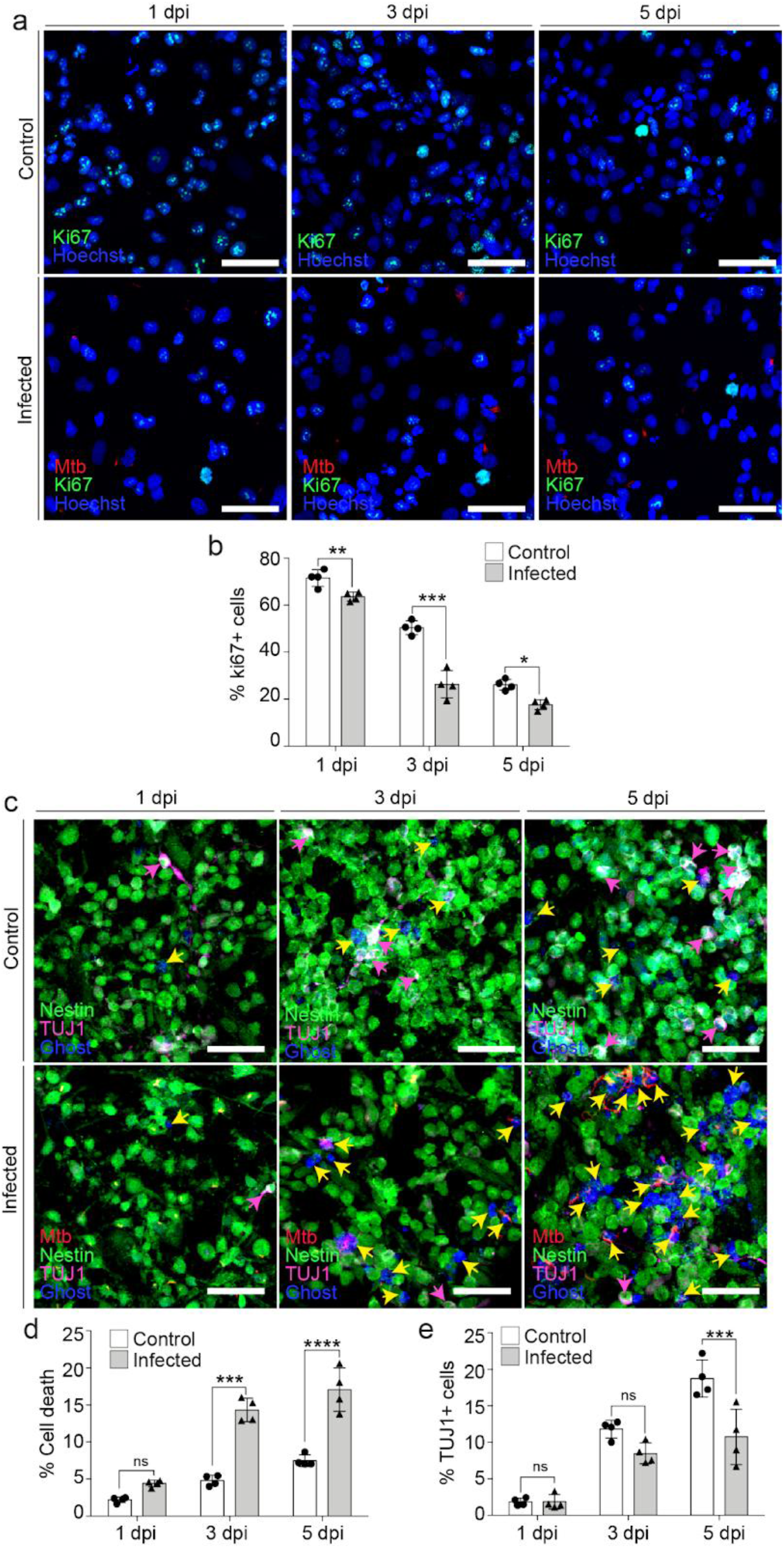
H37Rv Mtb-tdTomato decreases NPC proliferation and differentiation, and increases cell death. NPCs derived from hiPSCs were infected with H37Rv Mtb-tdTomato for 1, 3, or 5 days. Cells were stained for Ki67, cell viability dye (ghost), nestin, or beta III tubulin (TUJ1) and quantified using confocal microscopy. **(a)** Representative confocal images of uninfected and Mtb-infected NPCs stained for Ki67 (scale bars: 50 µm). **(b)** Frequencies of Ki67-positive cells in uninfected and infected cultures quantified by confocal microscopy. **(c)** Representative confocal images of uninfected and Mtb-infected NPCs stained for cell viability dye (ghost), nestin, and TUJ1 (scale bars: 50 µm). Yellow and magenta arrows indicate dead and differentiated cells, respectively. Frequencies of cell death **(d)** and TUJ1-positive **(e)** cells are presented in bar graphs. Data are shown as mean ± standard deviation (s.d.). Statistical significance: *p<0.05, ***p<0.001, ****p<0.0001.

### H37Rv Mtb infection induces type I IFN expression, and IFN-alpha neutralization antibodies (nAbs) partially mitigate Mtb-induced proliferative defects and cell death

Previous RNA sequencing analysis of H37Rv Mtb-infected hiPSC-derived NPCs revealed upregulation of gene signatures and biological pathways associated with type I IFN signaling, production, and responses. Type I IFNs are recognized regulators of cellular functions. To determine the expression of IFN-alpha (IFN-α) following H37Rv Mtb infection in hiPSC-derived NPCs, IFN-α expression was measured after infection using immunostaining followed by confocal microscopy. Induced IFN-α expression was observed in H37Rv Mtb-infected hiPSC-derived NPCs at 3 dpi compared to uninfected controls (Fig. 7a and b). Elevated levels of IFN-α subtypes, such as IFN-α1, have also been reported in West Nile virus (WNV)-infected human neural stem cells (NSCs) after 3 dpi(Riccetti et al. 2020). Low levels of type I IFNs (IFNa and IFNb) are required to elicit antibacterial immunity during the early phase of bacterial infections(McNab et al. 2015). However, adverse effects of type I IFNs on the host have been documented in intracellular bacterial infections, including Mtb and *Listeria monocytogenes*(McNab et al. 2015)*. Notably, Zheng et al. (2014) demonstrated that treatment with* mouse IFN-α in cultured mouse hippocampal NSCs induced apoptosis and suppressed proliferation(Zheng et al. 2014). These findings suggest that the observed cell death and reduced proliferation of hiPSC-derived NPCs may result from IFN-α expression and its adverse effects following H37Rv Mtb infection. To investigate the impact of blocking IFN-α activity on Mtb-induced cell death and proliferation defects, hiPSC-derived NPCs were infected with H37Rv Mtb in the presence of IFN-α neutralization antibodies (nAbs). After 3 dpi, samples were stained for Ki67 and the cell death marker ghost dye, and analyzed by confocal microscopy. hiPSC-derived NPCs infected with H37Rv Mtb in the presence of IFN-α nAbs exhibited increased cell proliferation (Fig. 7c and e) and reduced cell death (Fig. 7d and f) compared to the group not treated with IFN-α nAbs. These results indicate that IFN-α nAb treatment can rescue H37Rv Mtb-induced proliferation defects and cell death in hiPSC-derived NPCs. Collectively, these findings demonstrate that increased IFN levels play a central role in Mtb-induced changes in NPC function.

**Fig. 7.**
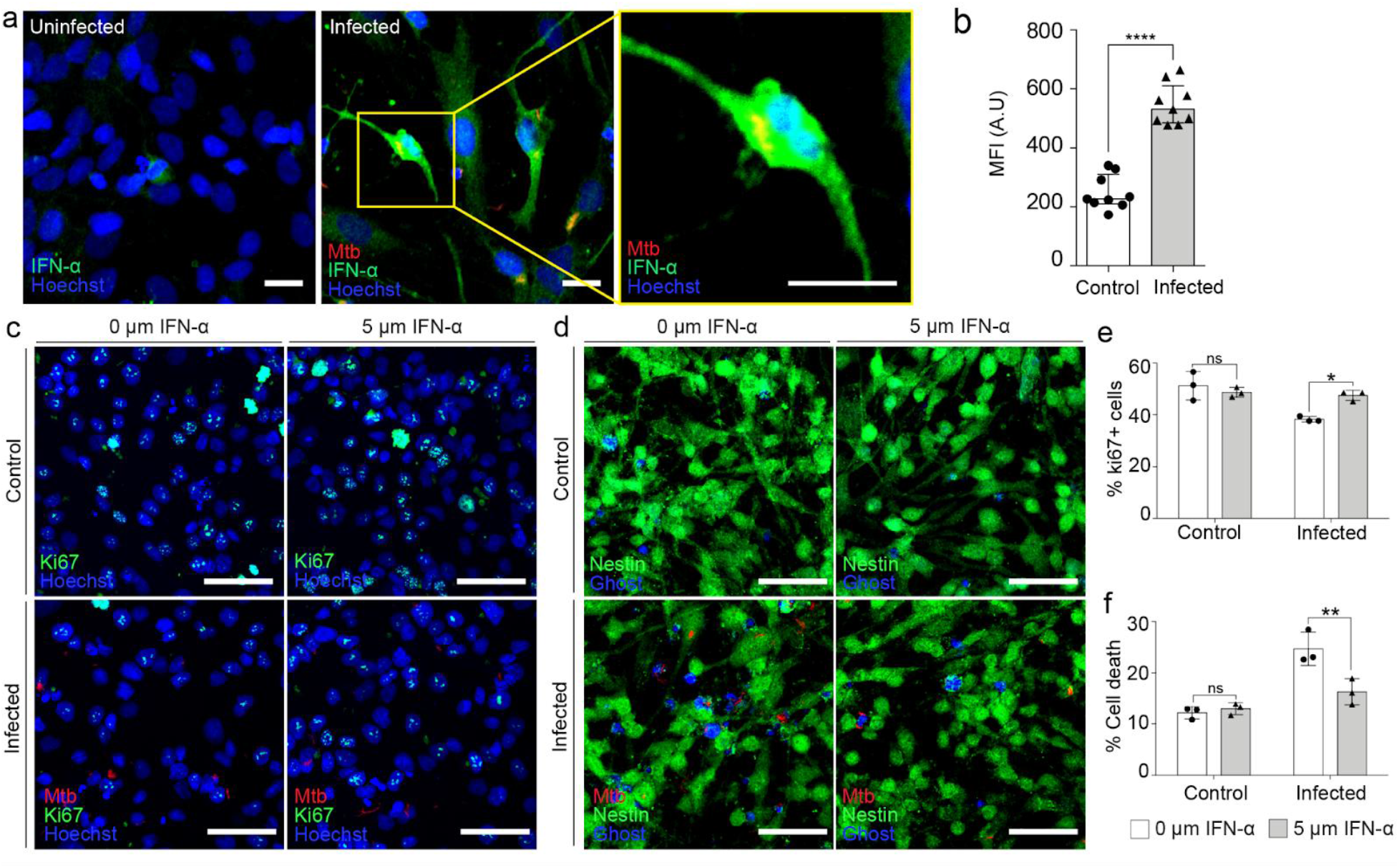
H37Rv Mtb-tdTomato infection promotes type I interferon expression. **(a)** Analysis of IFN-α expression in uninfected and Mtb-tdTomato (red) infected NPCs at 3 dpi using confocal microscopy. Scale bars: 10 µm. **(b)** Quantification of mean fluorescence intensity (MFI) of IFN-α in Mtb-infected and uninfected NPCs at 3 dpi. **Neutralizing antibodies against IFN-α (nAbs) partially inhibited Mtb-induced cell death in infected samples.** Uninfected and Mtb-infected NPCs were cultured for an additional 3 days in the presence or absence of 0.5 mM IFN-α nAbs. Proliferative and dead cells were stained with Ki67 and Ghost, respectively. **(c)** Representative confocal images of uninfected or Mtb-infected NPCs stained for Ki67 with or without IFN-α nAbs. **(d)** Representative confocal images of uninfected or Mtb-infected NPCs stained for dead cells with or without IFN-α nAbs. Frequencies of Ki67-positive **(e)** and dead cells **(f)** are presented in bar graphs. Data are shown as mean ± standard deviation (s.d.). (*p<0.05, **p<0.01).

### Mtb can be disseminated to NPCs from infected microglia

Because NPCs are localized in specific regions of the adult brain, systemic access of Mtb to NPCs may be limited. It was hypothesized that microglia, the resident macrophages of the CNS, could internalize Mtb and subsequently transmit the infection to NPCs. To investigate this mechanism, hiPSC-derived microglia were infected with tdTomato-expressing H37Rv Mtb for one hour at a multiplicity of infection (MOI) of 1:1, followed by staining with carboxyfluorescein succinimidyl ester (CFSE). The infected microglia were then co-cultured with hiPSC-derived NPCs and 3D hiPSC-derived neural organoids for one day. Interactions between H37Rv Mtb-infected microglia and hiPSC-derived NPCs were observed in 2D culture (Fig. 8a). Confocal orthogonal projections confirmed the internalization of H37Rv Mtb by hiPSC-derived NPCs in the 2D culture (Fig. 8b). Figure 8c outlines the experimental procedure for the co-culture system involving Mtb-infected microglia and 3D neural organoids. Comparable results were obtained in the co-culture system of infected microglia and 3D neural organoids. As shown in Figures 8d and 8e, H37Rv Mtb-infected nestin-positive NPCs were detected in 3D neural organoids co-cultured with infected microglia. These findings indicate that Mtb can be transmitted from infected microglia to NPCs.

**Fig. 8.**
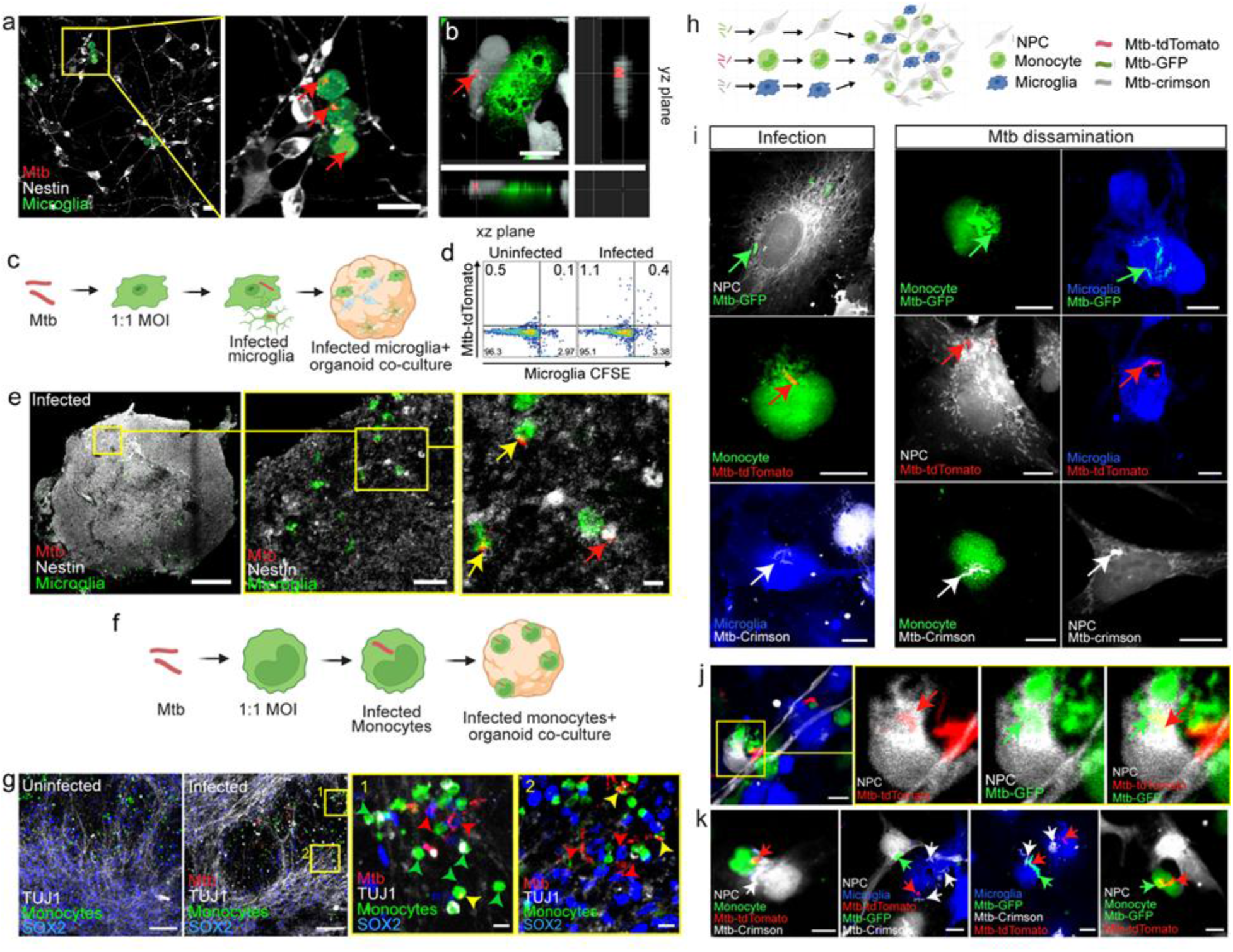
Mtb disseminates to NPCs in 2D cultures and 3D human neural organoids from infected microglia or monocytes. **(a-b)** hiPSC-derived microglia were infected with Mtb-H37Rv-tdTomato (red) at a multiplicity of infection of 1:1. After 1 hour, infected microglia were stained with CSFE and co-cultured with hiPSC-derived NPCs or human neural organoids for 1 day. **(a)** Representative confocal microscopy images illustrate the interaction between Mtb-infected microglia (green) and NPCs (white). **(b)** Orthogonal projections of confocal z-stacks (xy, yz, and xz planes) demonstrate Mtb dissemination from microglia into NPCs. Scale bars = 10 µm. **(c-e) Mtb disseminates to NPCs within neural organoids from infected microglia. (c)** Schematic representation of the experimental design. **(d)** Representative flow cytometry plots of human neural organoids co-cultured with either Mtb-infected or uninfected microglia; numbers indicate the percentage of infected cells. (e) Confocal microscopy images show Mtb-infected Nestin+ NPCs in human neural organoids co-cultured with infected microglia. Infected NPCs and microglia are indicated by red and yellow arrowheads, respectively. Scale bars = 100 µm; zoom: 10 µm. (**f-g) Mtb also disseminates to NPCs in neural organoids from infected human blood monocytes.** Human blood monocytes were infected with Mtb-H37Rv-tdTomato for 1 hour and stained with ViaFluor® 488 SE (green). Infected monocytes were then co-cultured with hiPSC-derived neural organoids for 1 day, followed by fixation and staining for SOX2 and TUJ1. **(f)** Schematic representation of the experimental setup. **(g)** Confocal microscopy images show neural organoids co-cultured with either uninfected (leftmost panel) or Mtb-infected human blood monocytes. Yellow boxes in the infected panel correspond to the two right panels, which display zoomed regions numbered 1 and 2. Infected NPCs and monocytes are indicated by red and yellow arrowheads, respectively. Scale bars = 100 µm; zoom: 10 µm. **(h-k) Mtb disseminates between hiPSC-derived microglia, NPCs, and human monocytes.** Human peripheral blood monocytes were infected with H37Rv Mtb-tdTomato at a multiplicity of infection of 1:1 for 1 hour, then labeled with ViaFluor® 488 SE (green). Infected monocytes were co-cultured with Mtb-crimson (grey) infected ViaFluor® 405-labeled microglia (blue) and Mtb-GFP-OVA (green) infected CellTrace™ Far Red-labeled NPCs (grey) for 1 day. **(h)** Schematic representation of Mtb dissemination among monocytes, microglia, and NPCs. (**i)** Confocal microscopy images show direct infection of Mtb in each phagocyte (left panel) and Mtb dissemination among monocytes, microglia, and NPCs in the co-culture system (right panel). Scale bars = 10 µm. Green, red, and white arrows represent Mtb-GFP, Mtb-tdTomato, and Mtb-Crimson, respectively. **(j-k) In the co-culture system, multiple bacteria can disseminate between individual phagocytes. (j)** A representative confocal image shows a single NPC infected with both Mtb-tdTomato and Mtb-GFP. **(k)** Representative confocal microscopy images demonstrate cross-dissemination of multiple bacteria into NPCs, microglia, and monocytes. Green, red, and white arrows indicate Mtb-GFP, Mtb-tdTomato, and Mtb-Crimson, respectively. Scale bars = 10 µm.

### Monocytes disseminate Mtb into other CNS phagocytic cells

Next, we investigated how systemic Mtb infects cells in the CNS. Previous studies suggest that systemic Mtb may cross the blood-brain barrier (BBB) via infected phagocytic cells, such as monocytes, subsequently disseminating infection to CNS cells(van Leeuwen et al. 2018; Wang et al. 2016). This process, termed the “Trojan horse mechanism,” has also been used to describe Zika virus transmigration through the BBB and subsequent infection of cerebral organoid cells(Ayala-Nunez et al. 2019). We have observed that infected monocytes and dendritic cells can traverse an in vitro BBB model(Gilpin et al. 2021). In our current experiment, we aimed to clarify whether Mtb infected monocytes can disseminate infection to CNS cells. As shown in Fig. 8f, human monocytes were infected with H37Rv Mtb-tdTomato at a MOI of 1:1 for 1 hour. The infected cells were then stained with ViaFluor® 488 SE and co-cultured with hiPSC-derived 3D neural organoids for 1 day. Confocal microscopy confirmed that infected blood monocytes interact with and disseminate Mtb to SOX2+ NPCs within the 3D neural organoids (Fig. 8g, two right-most panels).

We next sought to determine the potential for Mtb dissemination between different phagocyte populations within the co-culture system. Human monocytes, which are primary phagocytes potentially involved in trafficking systemic Mtb into the CNS, microglia (resident CNS phagocytes), and NPCs also displays Mtb phagocytic activity were infected with Mtb strains labeled with different color of fluorescent labels with H37Rv Mtb-tdTomato, H37Rv Mtb-Crimson, and H37Rv Mtb-GFP-OVA, respectively (Fig. 8h). Following infection, monocytes, microglia, and NPCs were stained with ViaFluor® 488 SE, ViaFluor® 405 SE, and Cell Trace Far Red, respectively, and co-cultured for 1 day (Fig. 8h). Confocal microscopy analysis demonstrated uptake of H37Rv Mtb by each cell type after direct infection (Fig. 8i, leftmost panel). Notably, we observed the dissemination of single bacilli of each fluorescently labeled H37Rv Mtb strain between the three cell types in the co-culture system (Fig. 8i, leftmost panel). For example, H37Rv Mtb-GFP OVA (green) primarily infected hiPSC-derived NPCs but was also detected in monocytes and microglia (Fig. 8i, upper panel). Furthermore, dissemination of multiple color-coded Mtb strains was observed in each cell type (Fig. 8j and k). These findings support the hypothesis that Mtb may access the CNS via infected blood monocytes and that these cells can cross-infect CNS cells with phagocytic activity, such as microglia and NPCs.

## Discussion

The precise identification of CNS cells targeted by Mtb invasion and the mechanisms facilitating bacterial uptake remain largely unresolved. While microglia are established as potential targets for Mtb(Spanos et al. 2015), the pathogen’s capacity to infect both professional(Maphasa et al. 2020; Lavalett et al. 2020; de Martino et al. 2019) and non-professional phagocytic cells(Mishra et al. 2023; Lerner et al. 2020; Khan et al. 2017) indicates the presence of additional cellular targets within the CNS. Previous studies have demonstrated that neurons(Randall et al. 2014) and astrocytes(Rock et al. 2005) are susceptible to Mtb infection. The complexity of Mtb interactions with diverse cell types highlights the necessity for a comprehensive investigation into potential target cells and the mechanisms underlying Mtb invasion in the CNS.

This study provides the first evidence that hiPSC-derived NPCs are capable of internalizing Mtb. The investigation comprehensively examines the mechanisms by which NPCs internalize Mtb and the subsequent effects on NPC functions, including proliferation, cell death, differentiation, type-1 IFN responses, and the potential intracellular fate of ingested Mtb. The initial detection of H37Rv Mtb-infected NPC subpopulations in neural organoids introduces a novel aspect to the spectrum of CNS cell types targeted by Mtb. Additionally, confirmation of Mtb uptake by both hiPSC-derived and ASC5007 (ATCC) NPCs in 2D culture systems underscores the significance of this unexpected phagocytic role. Previous in vivo and in vitro studies have documented the phagocytic activities of NPCs(Leeson et al. 2018; Lovelace et al. 2015; Ginisty et al. 2015; Lu et al. 2011). Notably, NPCs within the SVZ have demonstrated the ability to internalize PtdSer-containing liposomes and apoptotic NPCs in vivo(Leeson et al. 2018). In vitro, SVZ-derived neural stem-like cells or NPCs exhibit discernible phagocytic activity toward fluorescent latex beads or apoptotic neurons(Lovelace et al. 2015; Ginisty et al. 2015). These findings are consistent with the current observations. The loss of Mtb phagocytic activity in NPCs was observed upon their differentiation into neurons. Furthermore, experiments with various human clinical Mtb isolates and the Erdman strain demonstrated that all tested Mtb strains were internalized by hiPSC-derived NPCs at varying frequencies.

In vitro experiments with hiPSC-derived NPCs demonstrate that Mtb uptake occurs through phagocytic mechanisms involving membrane-bound receptors. The application of various phagocytosis inhibitors, such as annexin V and the P2X7R agonist BzATP, provides insight into the receptors potentially mediating Mtb internalization. In neurogenic niches, the uptake of dying cells by phagocytic cells is primarily mediated by recognition of PtdSer molecules on the cell surface, commonly referred to as the ‘eat me signal’(Gregory and Pound 2011; Elliott and Ravichandran 2010). PtdSer molecules on apoptotic cells are typically recognized directly by phagocyte surface receptors such as Bai1, Tim4, and Stabilin2, or indirectly through interactions with soluble factors that subsequently bind to other phagocytic receptors, including Mer and αvβ3/5(Elliott and Ravichandran 2010). Lu et al. demonstrated that Engulfment and cell motility protein 1 (ELMO1) is essential for regulating NPC phagocytic activity(Lu et al. 2011). The interaction between ELMO1 and the cytoplasmic tail of Bai1 is necessary to initiate cytoskeletal rearrangement via Dock180/Rac, facilitating the uptake of apoptotic cells(Park et al. 2007). In the present study, partial inhibition of H37Rv Mtb uptake by hiPSC-derived NPCs in the presence of annexin V suggests that NPCs may employ an efferocytosis-like process to internalize Mtb, potentially through recognition of PtdSer molecules by Bai1, PROS1 (protein S), and MerTK receptors. Additionally, P2X7R may participate in Mtb uptake when not engaged with ATP. The effects of cytoskeletal rearrangement inhibitors further underscore the active involvement of NPCs in this process.

The confirmation of phagolysosome colocalization of internalized H37Rv Mtb bacilli provides further evidence for the phagocytic activity of NPCs. This observation is supported by RNA sequencing data, which show enrichment of gene ontology cellular component (GO-CC) terms related to the phagocytic vesicle membrane and induction of genes such as *HLA-E, TAP1*, and *TAP2,* which are characteristic features of Mtb phagosomes(Harriff et al. 2013). However, the limited association of Mtb bacilli with membrane-bound endosomes raises the possibility that internalized Mtb may impede phagosomal maturation, potentially enabling escape from the phagosome. It is well established that internalized Mtb inhibits phagosome-lysosome fusion to promote bacterial survival in professional phagocytes such as macrophages (Jamwal et al. 2016). Mtb achieves this by manipulating host signaling pathways, resulting in retention within early endosomal compartments and eventual escape to the cytoplasm(Queval et al. 2017; Jamwal et al. 2016; Meena and Rajni 2010). However, the mechanisms of phagosomal escape in Mtb-infected non-myeloid cells remain poorly characterized. Khan et al. (2017) demonstrated that Mtb-infected mesenchymal stem cells limit intracellular bacterial growth through autophagy(Khan et al. 2017). While the survival and replication of intracellular pathogens such as Mtb in professional phagocytic cells are well documented, there is limited information regarding these processes in non-professional phagocytic cells. Mtb has been reported to infect, replicate, and persist in various non-professional phagocytes, including fibroblasts(Ferrer et al. 2009), HeLa cells(Shepard 1955), and lymphatic endothelial cells (LECs)(Lerner et al. 2020). This phenomenon highlights the need for a comprehensive understanding of Mtb dynamics in diverse cell types. Consistent with reports on Mtb-infected LECs(Lerner et al. 2020) and alveolar epithelial cells(Mishra et al. 2023), the present findings reveal the formation of intracellular cord-like structures in hiPSC-derived NPCs. Such cord formation enables bacteria to evade host immune responses and facilitates intercellular penetration, promoting dissemination into new tissue niches(Mishra et al. 2023). Ultimately, intracellular cord formation may contribute to Mtb survival within NPCs and offers novel insights into Mtb interactions with non-myeloid cells. Further investigation of these mechanisms may identify potential targets for interventions to disrupt Mtb survival across diverse host cell types.

In vitro analysis of H37Rv Mtb infection in hiPSC-derived NPCs demonstrates a marked reduction in Ki67-positive cells over time, indicating Mtb-induced defects in cellular proliferation. Additionally, increased cell death at later stages of infection suggests the involvement of factors such as intracellular Mtb growth or cord formation. The presence of uninfected dead cells adjacent to H37Rv Mtb-infected cells implies a possible mechanism by which infected cells induce cell death in neighboring cells. Previous studies have shown that infectious agents, such as the ZIKA virus, preferentially target NPCs and disrupt their differentiation(Li et al. 2016). Similarly, defective NPC differentiation was observed in H37Rv Mtb-infected samples, suggesting that Mtb-induced microenvironmental changes may impair NPC development. This is further supported by the downregulation of *NEUROD1*, *NEUROD4*, *NEUROG1,* and *NEUROG2*, genes involved in neuronal fate determination, terminal differentiation, and neurogenesis, which reinforces the evidence for Mtb-induced functional defects in hiPSC-derived NPCs(Wang et al. 2023; Fukuoka et al. 2021; Han et al. 2018; Lacomme et al. 2012; Boutin et al. 2010). Collectively, these findings warrant further investigation into whether Mtb can infect NPCs in adults and the potential impact on adult neurogenesis.

RNA sequencing data from H37Rv Mtb-infected hiPSC-derived NPCs revealed enrichment of pathways associated with type-1 IFN signaling and upregulation of key type-1 IFN-responsive genes, as well as genes involved in type-1 IFN production. While type-1 IFN responses are typically prominent during viral infections, Mtb-infected macrophages(Andreu et al. 2017), THP cells(Wu et al. 2012), and non-myeloid cells such as human lymphatic endothelial cells (LECs)(Lerner et al. 2020) and human retinal pigment epithelial cells (RPE)(La Distia Nora et al. 2018) also exhibit significant expression of type-1 IFN-stimulated genes, consistent with the present findings. Supporting this, single-cell RNA sequencing of attenuated *Mycobacterium bovis*-infected mouse brains demonstrated enrichment of IFN-α response terms in NPCs(Zhang et al. 2023). Beyond cellular consequences, the induction of type-1 IFN expression in NPCs following Mtb infection may have detrimental effects. Previous research has linked IFN-α to inhibition of NPC proliferation, reduced neurogenesis, and initiation of apoptosis in various experimental models(Borsini et al. 2018; Zheng et al. 2014). The current results indicate that activation of type-1 IFNs in NPCs after H37Rv Mtb infection may result in decreased proliferation, increased cell death, and impaired neurogenesis. These changes in neural processes may influence cognitive behavior, as prior studies have suggested associations between type-1 IFNs, hippocampal neurogenesis, and depressive phenotypes. Notably, treatment with IFN-α neutralizing antibodies partially rescued hiPSC-derived NPCs from Mtb-induced cell death and proliferative defects, suggesting that type-1 IFNs produced during Mtb infection play a detrimental role in NPCs. IFN-α neutralizing antibody treatment may represent a potential therapeutic strategy for mitigating CNS-TB-induced cellular complications, although further studies are necessary to confirm this approach.

NPCs are primarily located in the SVZ and hippocampal SGZ, which serve as neurogenic niches(Alvarez-Buylla and Garcia-Verdugo 2002; Eriksson et al. 1998), and typically have limited exposure to Mtb. Mtb is found in phagocytes, including blood-derived monocytes recruited from the periphery, and in microglia, the resident macrophages of the brain responsible for phagocytic clearance. In vivo, NPCs likely become infected with Mtb through contact with infected monocytes and microglia, which may serve as carriers. Supporting this, H37Rv Mtb-infected NPCs were identified in 2D NPC plus H37Rv Mtb-infected microglia cultures, 3D neural organoid plus H37Rv Mtb-infected microglia, and 3D neural organoid plus H37Rv Mtb-infected monocyte co-culture systems, indicating possible dissemination of Mtb from microglia and monocytes to NPCs. Furthermore, the observation of distinct color-coded H37Rv Mtb cross-dissemination among microglia, NPCs, and monocytes in co-culture systems suggests that peripheral Mtb may employ a “trojan horse-like mechanism” to traverse the blood-brain barrier and infect NPCs within the CNS.

In summary, this research advances the understanding of Mtb interactions with NPCs in the CNS. The identification of an unexpected phagocytic role for NPCs, along with observed effects on neurogenesis, cell death, and immune responses, highlights the complex consequences of Mtb infection in the brain. These findings underscore the need for further investigation into the molecular mechanisms involved and the development of potential therapeutic interventions.

## Materials and Methods

### Cells

H1 human embryonic stem cells (ES) derived NPCs (hiPSC-derived NPC) and microglia were obtained from Dr. J. Thompson (University of Wisconsin-Madison, Madison, WI, USA). hiPSC derived NPCs and Neural Progenitor Cells Derived from XCL-1 MAP2p-Nanoluc-Halotag (ASC-5007, ATCC) were cultured in Matrigel (growth factor reduced, Catalog No:356230, Corning) coated plates in DF3S (DMEM/F-12 (Catalog No: 11330032, Gibco), sodium selenite (14 μg/L; Catalog No: S5261, Sigma-Aldrich) and and NaHCO3 (543 mg/L) media supplemented with 1x N2 (Catalog No: 17502048, Gibco), 2% B27 (Catalog No: 17504044, Gibco) and recombinant human FGF2 (rhFGF2) (5 μg/L, Catalog No: SRP4037-50UG, Sigma-aldrich)(Schwartz et al. 2015). hiPSC-derived microglia were maintained in IMDM (Gibco) supplemented with 10% FBS, rhM-CSF (20 ng/mL) (Peprotech) and rhIL-1β (10 ng/mL) (Peprotech)(Schwartz et al. 2015). Purified human monocytes that were derived from peripheral blood mononuclear cells (PBMCs) were purchased from iQ BIOSCIENCES (Catalog No: IQB-Hu1-M10) and cultured according to the manufacturer’s instructions. All cells were grown in antibiotic free media and maintained at 37°C and 5% CO_2_ in a humidified incubator.

### Mtb Strains and Culture

*Mycobacterium tuberculosis* (Mtb) H37Rv was obtained from Dr. Adel Talaat (University of Wisconsin-Madison, WI). The Mtb strain was engineered with tdTomato (pTEC27) plasmids (Catalog No: 30182, Addgene), as described by Takaki et al(Takaki et al. 2013), as well as Crimson (pTEC19) and GFP-OVA. Strains were stored as frozen aliquots at - 80 °C. The Mtb Erdman strain was provided by Dr. Shelby O’Connor (University of Wisconsin-Madison). Human clinical Mtb isolates, including brain (SK81) and sputa (SK131 and SK139), were obtained from Dr. Caitlin Pepperell (University of Wisconsin-Madison). All Mtb strains were cultured at 37 °C in Middlebrook 7H9 medium (Catalog No: 271310, BD) supplemented with 10% oleic acid-albumin-dextrose-catalase (OADC) enrichment (Catalog No: B11886, BD) and 0.05% Tween 80 (Catalog No: P4780, Sigma Aldrich), with or without hygromycin B (100 μg/mL; Catalog No: ant-hm-1, InvivoGen), in a shaking incubator or on Middlebrook 7H10 agar medium (BD) with 10% OADC and 0.5% glycerol (Catalog No: BP2291, Fisher Scientific) at 37 °C. All procedures involving Mtb growth, infection, sample processing, and collection were performed in a Biosafety Level 3 (BSL3) facility in accordance with approved biosafety guidelines.

### Mtb infection of NPCs

hiPSC-derived NPCs or XCL-1 MAP2p-Nanoluc-Halotag (ASC-5007, ATCC) NPCs were seeded as a monolayer in Matrigel-coated, glass-bottom 8-well Nunc Lab-Tek Chamber Slide Systems (Catalog No: 177402PK, Thermo Fisher Scientific) at a density of 4 × 10^4^ cells per well in NPC complete medium and allowed to adhere overnight. The following day, cells were infected with H37Rv, Erdman, or Mtb clinical isolates at a multiplicity of infection (MOI) of 1:1 for 1, 3, or 5 days. At 1 day post-infection (1 dpi), cells were washed thoroughly with pre-warmed sterile 1× Dulbecco’s Phosphate-Buffered Saline (DPBS) (Catalog No: 14190144, Gibco), and fresh pre-warmed medium was added. Half of the medium was replaced every other day until the conclusion of the experiment.

### Human Neural Organoids

Human neural organoids were obtained from StemPharm, Inc. (Madison, WI) in a 96-well format. Organoids were formed on polyethylene glycol-based synthetic hydrogels as previously described and maintained according to the manufacturer’s recommendations under standard cell culture conditions, with half-media changes performed daily(Barry et al. 2017; Schwartz et al. 2015). Human NPCs derived from XCL-1 and obtained from ATCC (ACS-5007)(Pei et al. 2016, 2015) were used to generate the neural organoids. XCL-1 is a subclone derived from NCRM1, an integration-free iPSC line (NIH Center for Regenerative Medicine) established from CD34+ cord blood of a newborn male. NPCs were cultured in STEMdiff™ Neural Progenitor Medium (NPM) (Stem Cell Technologies) and expanded on a thin coating of Geltrex (Thermo; 1:100 dilution in DMEM/F12, Thermo). Prior to experimentation, cells were cultured for at least one passage on Geltrex. Neural organoid cultures were utilized in experiments at 30 days post-plating. Thirty-day-old neural organoids were infected with H37Rv Mtb-tdTomato (colony-forming units [CFU]=1×10^5^) for three days. Subsequently, the neural organoids were thoroughly washed with PBS and fixed with 4% paraformaldehyde (PFA) (Catalog No: 15714-S, Electron Microscopy Sciences) for one day.

### In vitro NPC Viability Assay

Dead cells were stained using Ghost Dye™ Violet 450 (SKU 13-0863-T100, Cytek). Uninfected or H37Rv Mtb-infected NPCs cultured as monolayers in 8-well chamber slides were collected at 1, 3, and 5 days dpi. Cells were washed thoroughly with 1× PBS, and the medium was replaced with dead cell staining dye (1:1000) in PBS. Cells were incubated for 20 minutes at room temperature, protected from light. Subsequently, cells were washed once with PBS and fixed with 4% PFA for 1 hour. PFA was replaced with PBS, and cells were stored at 4 °C until further processing.

### Mtb Uptake Inhibition Assay

hiPSC-derived NPCs were cultured in Matrigel-coated 6-well tissue culture plates (Catalog No: 229105, CELLTREAT) or 8-well glass chamber slides at a density of 5 × 10^5^ or 4 × 10^4^ cells per well, respectively, in complete NPC medium overnight prior to experimentation. Cells were infected with H37Rv Mtb-tdTomato at a MOI of 1:1, in the presence or absence of 0.68 mM purified annexin V (Catalog No: 640901, BioLegend), 30 mM cytochalasin D (Catalog No: C273-1MG, Sigma), 250 mM 2′(3′)-O-(4-Benzoylbenzoyl)adenosine 5′-triphosphate triethylammonium salt (BzATP) (Catalog No: B6396-5MG, Sigma-Aldrich), and 25 mM blebbistatin (Catalog No: B0560-1MG, Sigma-Aldrich) for 1 day. Following infection, cells in 6-well plates were washed with PBS, dissociated using StemPro™ Accutase™ Cell Dissociation Reagent (Catalog No: A1110501, Gibco) and mechanical disruption, centrifuged, washed again with PBS, and resuspended in 4% PFA for 1 hour. Subsequently, cells were washed once with PBS and resuspended in FACS buffer (1% bovine serum albumin in PBS) for flow cytometric analysis. Cells in 8-well chamber slides were washed with PBS, fixed with 4% PFA for 1 hour, and then maintained in PBS at 4°C until further immunocytochemical analysis.

### hiPSC-Derived NPC/Neural Organoid and Microglia Co-Culture System

hiPSC-derived microglia were seeded in 6-well plates at a density of 2 × 10^5^ cells per well in DM5 medium and incubated at 37°C overnight. On the day of infection, cells were exposed to H37Rv Mtb-tdTomato at an MOI of 1:1 for 1 hour. Following infection, cells were washed with 1X PBS and stained with CFSE (Catalog No: 13-0850-U500, Tonbo Biosciences) according to the manufacturer’s instructions. After staining, hiPSC-derived microglia were dissociated using StemPro™ Accutase™ Cell Dissociation Reagent, and viable cell counts were determined. Subsequently, 1 × 10^4^ microglia were co-cultured with 4 × 10^4^ uninfected hiPSC-derived NPCs in Matrigel-coated 8-well chamber slides in NEM for 1 day. In parallel, 2 × 10^4^ hiPSC-derived microglia were co-cultured with neural organoids in hNMM for 1 day. After 1 day post-infection, both co-culture systems were fixed with 4% PFA and processed for immunocytochemical analysis.

### hiPSC-Derived Neural Organoids and Monocytes Co-Culture System

Human monocytes were cultured in 6-well plates at a density of 2 × 10^5^ cells per well in RPMI 1640 (Catalog No: 10-040-CV, Corning) supplemented with 10% fetal bovine serum. Cells were then infected with H37Rv Mtb-tdTomato at an MOI of 1:1 for 1 hour. Following infection, cells were washed with 1X PBS to remove excess bacteria and stained with ViaFluor® 488 SE (Catalog No: 30086-T, Biotium) according to the manufacturer’s instructions. A total of 2 × 10^4^ stained monocytes were co-cultured with human neural organoids in hNMM for 1 day. After 1 day post-infection, the co-culture system was fixed with 4% PFA and processed for immunocytochemical analysis.

### hiPSC-Derived NPC, Microglia, and Purified Human Monocyte Co-Culture System

hiPSC-derived NPCs were infected with H37Rv Mtb-GFP-OVA at an MOI of 1:1 for 1 day, followed by staining with CellTrace™ Far Red (Catalog No: C34572, Invitrogen) according to the manufacturer’s instructions. Similarly, hiPSC-derived microglia and purified human monocytes were infected with H37Rv Mtb-crimson and H37Rv Mtb-tdTomato, respectively, at an MOI of 1:1 for 1 hour. Infected hiPSC-derived microglia and human monocytes were then stained with ViaFluor® 405 SE (Catalog No: 30068-T, Biotium) and ViaFluor® 488 SE (Catalog No: 30086-T, Biotium), respectively. The three cell types H37Rv Mtb-GFP-OVA-infected hiPSC-derived NPCs (stained with CellTrace™ Far Red), H37Rv Mtb-crimson-infected hiPSC-derived microglia (stained with ViaFluor® 405 SE), and H37Rv Mtb-tdTomato-infected human monocytes (stained with ViaFluor® 488 SE) were co-cultured in NEM in Matrigel-coated 8-well chamber slides for 1 day. Following co-culture, cells were fixed with 4% PFA and processed for immunocytochemical analysis.

### IFN-α Neutralization

hiPSC-derived NPCs were seeded as a monolayer at a density of 4 × 10^4^ cells per well in Matrigel-coated 8-well glass-bottom Nunc Lab-Tek Chamber Slides, using NPC complete medium. Cells were allowed to adhere overnight. Subsequently, cells were infected with H37Rv Mtb-tdTomato at a MOI of 1:1 for either 1 or 3 days in the presence of human IFN-α neutralizing antibody (5 μg/mL, MMHA-17, Catalog No: 21118-1, PBL Biomedical Laboratories). At designated time points, samples were fixed with 4% PFA for 1 hour and processed for immunocytochemistry analysis.

### Lysotracker Staining

At 1 dpi, H37Rv Mtb-tdTomato-infected hiPSC-derived NPCs were washed with 1X PBS and incubated with 75 nM LysoTracker Deep Red (Catalog No: L12492, Invitrogen) in pre-warmed NEM at 37 °C for 1 hour. Following incubation, cells were washed twice with PBS, fixed with 4% PFA for 1 hour, and processed for immunocytochemistry analysis.

### Auramine O Staining

Auramine O staining was used to fluorescently label three clinical Mtb isolates and the Mtb-Erdman strain. Briefly, Mtb-infected NPCs fixed with 4% PFA were washed three times with 1X PBS, then permeabilized with 0.1% Triton X-100 (Catalog No: H5141), Promega) in PBS for 30 minutes. Cells were treated with freshly prepared auramine O solution (0.1% Auramine [Catalog No: A968-25, Fisher Scientific] and 3% Phenol [Catalog No: P1037-25G, Sigma-Aldrich]) for 20 minutes. After thorough washing with distilled water three times, cells were treated with acid alcohol solution (concentrated HCl in 70% ethanol) for 3 minutes. Cells were again washed three times with distilled water, incubated with 0.5% potassium permanganate (Catalog No: 223468-25G, Sigma) solution for 1 minute, and washed three more times with distilled water before air drying. Nuclei were counterstained using Hoechst 33342 (3 µg/mL, Catalog No: 382065, Sigma-Aldrich) and mounted with Prolong Glass antifade mountant (Catalog No: P36980, Invitrogen).

### Transmission Electron Microscopy (TEM)

hiPSC-derived NPCs were cultured on Matrigel-treated 12 mm round glass coverslips (Catalog No: CLS1760012, Chemglass Life Sciences) at a density of 2.5 × 10^5 cells per well in complete NPC medium overnight before the experiment. The cells were infected with H37Rv Mtb-tdTomato at an MOI of 1:1 for 1 and 3 days. At each time point, samples were fixed with 4% PFA for 1 hour, then replaced with 2% buffered glutaraldehyde solution. Samples were post-fixed with 2% osmium tetroxide (Electron Microscopy Sciences), dehydrated using graded ethanol solutions, and embedded in epoxy resin. Samples were sectioned using an ultramicrotome, and Post-staining was performed with 8% uranyl acetate in 50% methanol and lead citrate, followed by analysis using TEM (Philips CM 120) (Hirano et al. 2003).

### Flow Cytometry

To determine the frequency of Mtb-positive NPCs in infected groups and in the Mtb uptake inhibition experiment, cells resuspended in fluorescence-activated cell sorting (FACS) buffer (PBS + 1% BSA) were analyzed using a NL 3000 V/B/R spectral flow cytometer (Cytek Biosciences). Data were analyzed using FlowJo software (BD Biosciences).

### Immunostaining and Confocal Microscopy

Immunocytochemistry was performed on Mtb-infected and uninfected NPC monolayers. Cells were washed twice with PBS for 10 minutes each, followed by incubation with blocking and permeabilization buffer (PBS with 1.0% BSA, 0.25 % Triton X-100) for 1 hour at room temperature. Primary antibodies were applied in staining buffer (PBS with 1.0% BSA, 0.1% Triton X-100) and incubated at 4 °C overnight. The following primary antibodies were used: nestin (1:500, Catalog No: NB100-1604, Novus Biologicals), Tubulin b3 (TUJ1) conjugated to Alexa Fluor 647 (1:200, Catalog No: 801210, BioLegend), Ki67 (1:500, Catalog No: ab15580, Abcam), LAMP-1 (1:100, Catalog No: 328602, BioLegend), Rab5A (Catalog No: NBP1-04340SS, Novus Biologicals), Rab7A (Catalog No: NBP2-24591SS, Novus Biologicals), and IFN-α (MMHA-17) (1:200, Catalog No: 21118-1, PBL Biomedical Laboratories). Cells were then washed three times with PBS for 10 minutes each and incubated with secondary antibodies diluted in staining buffer, with or without the nuclear stain Hoechst 33342 (3 µg/mL, Catalog No: 382065, Sigma-Aldrich), at room temperature for 1 hour. The following secondary antibodies were used: goat anti-chicken Alexa Fluor 488 (1:500, Catalog No: A-11039, Invitrogen), goat anti-rabbit Alexa Fluor 488 (1:500, Catalog No: A-11008, Invitrogen), goat anti-mouse Alexa Fluor 488 (1:500, Catalog No: A11001, Invitrogen), and goat anti-rabbit Alexa Fluor 647 (Catalog No: A-21244, Invitrogen). After three additional PBS washes, cells were mounted with Prolong Glass antifade mountant (Catalog No: P36980, Invitrogen). Images were acquired using an Olympus FV1200 IX83 confocal microscope (Olympus, Shinjuku, Japan) and analyzed with IMARIS and ImageJ (National Institutes of Health open-source software).

For immunostaining of neural organoids, whole organoids were washed three times with PBS for 10 minutes each. Organoids were then permeabilized and blocked in a buffer containing 0.25% Triton X-100 and 5% BSA in PBS for 1 hour at room temperature. Primary antibodies, including nestin (1:500, Catalog No: NB100-1604, Novus Biologicals), SOX2, and Tubulin b3 (TUJ1) conjugated to Alexa Fluor 647 (1:200, Catalog No: 801210, BioLegend), were applied in staining buffer containing 0.1% Triton X-100 and 1% BSA in PBS and incubated overnight at 4 °C. Organoids were subsequently washed three times with 1X PBS for 10 minutes each. Secondary antibodies, including goat anti-chicken Alexa Fluor 488 (1:500, Catalog No: A-11039, Invitrogen) and Hoechst 33342 (3 µg/mL), were diluted in staining buffer and incubated with the organoids overnight at 4 °C. After three additional washes with 1X PBS for 10 minutes each, organoids were imaged using an Olympus FV1200 IX83 confocal microscope.

### Total RNA Extraction and Sequencing

H37Rv Mtb-tdTomato-infected and uninfected hiPSC-derived NPCs at 1 and 3 dpi were lysed in 1 ml of QIAzol Lysis Reagent (Catalog No: 79306, QIAGEN). Total RNA was extracted using the miRNeasy Micro Kit (Catalog No: 217084, QIAGEN) according to the manufacturer’s instructions. The quantity and quality of extracted RNA were assessed using a Nanodrop spectrophotometer (Thermofisher Scientific). Subsequent steps, including RNA integrity assessment with an Agilent Bioanalyzer, library construction, and next-generation sequencing on the Illumina platform, were performed by Active Motif (WWW.activemotif.com).

### RNA-seq Analysis

Quality assessment of raw paired-end FASTQ files was performed using FastQC software to ensure accurate base calling and the absence of sequencing abnormalities. The base content was confirmed to be unbiased throughout the reads. Following quality assessment, RSEM software was used to generate read counts. Bowtie2 was employed to align the reads to the human genome (hg38), achieving an average mapping efficiency of 88% across all samples. Gene count matrices were generated, and differential expression analysis was conducted using the R package EBSeq. Differentially expressed genes (DEGs) were identified using a posterior probability of differential expression greater than 0.95, corresponding to an adjusted P-value of less than 0.05, and a posterior log2 fold-change greater than or equal to one in absolute value. For Gene Ontology enrichment and pathway analysis, DEGs were analyzed using the SRplot(Tang et al. 2023) and Reactome pathway databases, respectively(Milacic et al. 2024).

### Statistical Analysis

Statistical analyses were conducted using GraphPad Prism 6.0 software. An unpaired Student’s t-test was used to compare results between two groups. Data are presented as mean ± standard deviation (SD), and statistical significance is indicated as *P < 0.05, **P < 0.01, ***P < 0.001, and ****P < 0.0001.

## Acknowledgment

This work was supported by National Institutes of Health (NIH) grant no. 1R56AI162164 and 5R01NS123449 (awarded to M.S.). C.L. was supported by AHA grant 915125 and in part by NIH/NINDS T32 NS105602.

